# Pathogenic mis-splicing of *CPEB4* in schizophrenia

**DOI:** 10.1101/2022.09.22.508890

**Authors:** Ivana Ollà, Antonio F. Pardiñas, Alberto Parras, Ivó H. Hernández, María Santos-Galindo, Sara Picó, Luis F. Callado, Ainara Elorza, Gonzalo Fernández-Miranda, Eulàlia Belloc, James T.R. Walters, Michael C. O’Donovan, Claudio Toma, Raúl Méndez, J. Javier Meana, Michael J. Owen, José J. Lucas

**Author notes:** To whom correspondence should be addressed: José J. Lucas, Center for Molecular Biology-Severo Ochoa (CBMSO), C/ Nicolás Cabrera,1. 28049 Madrid. Spain, http://www.cbm.uam.es/LucasLab.

## Abstract

Schizophrenia (SCZ) is caused by a complex interplay of polygenic risk and environmental factors, which might alter regulators of gene expression leading to pathogenic mis-expression of SCZ risk genes. The RNA binding protein family CPEB (CPEB1, CPEB2, CPEB3, CPEB4) regulates the translation of target RNAs bearing CPE sequences in their 3’UTR (approximately 40% of overall genes). We previously identified CPEB4 as a key dysregulated translational regulator in autism spectrum disorder (ASD), proving that its neuronal-specific microexon (exon 4) is mis-spliced in brains of ASD probands, leading to concerted underexpression of a plethora of high confidence ASD-risk genes. The genetic and pathogenic mechanisms shared between SCZ and ASD make it plausible that mis-splicing of CPEB4 might occur also in SCZ patients, leading to downstream altered brain expression of multiple SCZ-related genes. In this study, we first analyzed Psychiatric Genomics Consortium GWAS data and found significant enrichment of SCZ-associated genes for CPEB4-binder transcripts. We also found decreased inclusion of CPEB4 microexon in postmortem prefrontal cortex of SCZ probands. This mis-splicing is associated with decreased protein levels of SCZ-associated genes that are targets of CPEB4. Interestingly, this happens specifically in individuals with low exposure to antipsychotic medication. Finally, we show that mild overexpression of a CPEB4 transcript lacking exon 4 (CPEB4Δ4) in mice suffices to induce decreased protein levels of SCZ genes targeted by CPEB4; these mice are also characterized by SCZ-linked behaviors. In summary, this study identifies aberrant CPEB4 splicing and downstream mis-expression of SCZ-risk genes as a novel etiological mechanism in SCZ.

## Introduction

Schizophrenia (SCZ) is a severe psychiatric disorder characterized by abnormalities in thought and cognition which affects nearly 1% of the adult population (Owen et al., 2016). Genetic and epidemiological evidence indicate that SCZ is caused by common and rare risk alleles combined with environmental factors (Henriksen et al., 2017). In the last decade, advances in genomics and collaborative efforts from international consortia have identified around 300 risk alleles although biological and pathophysiological mechanisms are still largely unknown (Hall et al., 2015; PGC, 2015). The identified genetic risk variants range from rare copy number variants, multiple rare single nucleotide variants and loci containing common genetic variants, the latter exerting individually small effect sizes (Henriksen et al., 2017). Since the precise molecular determinants that integrate polygenic risk and environmental risk factors in SCZ are not fully elucidated, it is important to investigate altered regulators of gene expression in the brains of individuals with SCZ that could explain orchestrated pathogenic mis-expression of numerous risk genes during both neurodevelopment and adult life.

Cytoplasmic polyadenylation element binding proteins (CPEBs) are a family of RNA-binding proteins that regulate the stability and translation of specific mRNAs containing CPE sequences in their 3’ untranslated regions (UTRs) (Ivshina et al., 2014) accounting for about 40% of the transcriptome (Parras et al., 2018; Pique et al., 2008). In vertebrates, the family consist of four paralogs (CPEB1, CPEB2, CPEB3 and CPEB4) where CPEB2-4 are closely related and CPEB1 is the most distant member of the family (Wang and Cooper, 2010). CPEBs mediate translational repression or activation of their target transcripts by inducing, respectively, shortening or elongation of the poly(A)-tails (Ivshina et al., 2014). CPEBs were first discovered through their role in development, as they regulate the expression of multiple mRNAs in response to embryonic environmental clues, such as hormones (Ivshina et al., 2014; Sarkissian et al., 2004). More recently, CPEBs have been shown to play important roles in cell division and metabolism (Alexandrov et al., 2012; Burns and Richter, 2008; Ivshina et al., 2014). In the adult brain, CPEBs are known to regulate many genes involved in synaptic plasticity, thus contributing to complex processes such as learning and memory consolidation (Drisaldi et al., 2015; Fioriti et al., 2015; Ivshina et al., 2014; Pavlopoulos et al., 2011; Si et al., 2003; Wu et al., 1998). Consistently, altered CPEBs have been associated with cancer (Ortiz-Zapater et al., 2011; Perez-Guijarro et al., 2016) and pathogenic hepatic angiogenesis (Calderone et al., 2016), neurological diseases including epilepsy (Parras et al., 2020) and Huntington’s disease (Picó et al., 2021) as well as with ASD (Parras et al., 2018; Udagawa et al., 2013).

In ASD, CPEB4 has emerged as a key pathogenic effector that is altered in brain tissues of cases, resulting in the simultaneous mis-expression of most high confidence ASD-risk genes (Parras et al., 2018). More precisely, we reported that individuals with ASD show an imbalance of CPEB4 transcript isoforms resulting from a decreased inclusion of a neuronal-specific microexon (exon 4) (Parras et al., 2018). Microexons, which are exons of 27 or fewer nucleotides, show a pattern of neural specific alternative splicing (AS) that has been shown to be dysregulated globally in ASD (Irimia et al., 2014; Li et al., 2015). We have also shown that the resulting increase of CPEB4Δ4 transcript isoforms in ASD brain tissues correlates with decreased protein levels of multiple ASD-risk genes whose transcripts harbour CPE sequences in their 3’UTR (Parras et al., 2018). Furthermore, we found that transgenic mice overexpressing CPEB4Δ4 showed decreased protein expression of a plethora of ASD risk genes and display ASD-like traits (Parras et al., 2018).

Schizophrenia and ASD share genetic risk, and by inference pathogenic mechanisms, and have been proposed to lie on a neurodevelopmental continuum (Owen and O’Donovan, 2017). This led us to hypothesize that alteration of CPEBs, particularly CPEB4, might also be observed in the brains of individuals with SCZ, and that this would lead to pathogenic mis-expression of multiple SCZ-risk-genes.

## Results

### SCZ susceptibility loci show enrichment for CPE-harboring and CPEB4-binder transcripts

We decided to explore whether CPE-containing and CPEB4-binding transcripts were over-represented within SCZ-associated genes. We first used MAGMA to perform gene-set analyses based on the summary statistics from the Wave 3 Psychiatric Genomics Consortium (PGC) GWAS of schizophrenia (“core dataset”; 67,390 cases and 94,015 controls) (Trubetskoy et al., 2022). Successively, we examined gene sets comprised of (i) genes containing canonical CPE (cCPE) sequences in their 3’UTR (Pique et al., 2008); (ii) genes identified in genome-wide RNA immunoprecipitation analyses from mouse brain structures as CPEB1- (Parras et al., 2018), CPEB3- (Lu et al., 2021) or CPEB4- (Parras et al., 2018) binders; (iii) gene sets implicated in the pathophysiology of psychiatric disorders: Fragile X Mental Retardation Protein (FMRP) targets (Clifton et al., 2021; Darnell et al., 2011), genes specifically involved in synaptic development and function (Koopmans et al., 2019), and finally (iv) as a comparator, a more general set of all brain-expressed genes (Fagerberg et al., 2014). The results of associations with SCZ-associated genes showed “CPEB4 target” as the top significant gene-set amongst those under study (*P*=1,76×10^−9^; Table 1), together with already known gene-sets implicated in psychiatric disorders such as FMRP targets and brain-expressed genes (*P*=4.97×10^−9^ and *P*=6.13×10^−19^, respectively; Table 1). MAGMA models were conditioned on a binary indicator to control for overlapping genes amongst gene-sets, given the known substantial overlap between FMRP and CPEB targets (Udagawa et al., 2013), or CPEB target genes and brain expressed genes (Parras et al., 2018). After conditional analysis for each of FMRP targets, synaptic genes, and brain-expressed genes, the “CPEB4 target” gene-set remained the most significant (Table 1). As expected, given the overlap among the CPEB1-, CPEB3-, and CPEB4-targets, CPEB1 and CPEB3 targets also showed evidence for enrichment for SCZ associations as did the gene set of “cCPE-containing genes” (Table 1). These results therefore confirmed enrichment of CPE-containing and CPEB4-binding transcripts in SCZ susceptibility genes.

**Table 1:** Gene-level analysis performed with CPE-containing, CPEB-targets and psychiatric-related gene sets using the latest GWAS summary statistics from the PGC of schizophrenia.

|  |  |  | Conditional Analysis (*P*-value) |  |  |
| --- | --- | --- | --- | --- | --- |
| Gene-set name | N genes | *P*-value | Brain-expressed | Synaptic genes | FMRP targets |
| cCPE-containing genes | 7,712 | 2.92E-04 | 0.036 | 5.93E-04 | 0.003 |
| CPEB1 targets | 2,207 | 1.39E-07 | 2.08E-04 | 7.06E-07 | 3.71E-06 |
| CPEB3 targets | 3,508 | 6.95E-04 | 0.065 | 0.002 | 0.003 |
| CPEB4 targets | 2,749 | 1.76E-09 | 5.77E-05 | 2.19E-08 | 7.98E-08 |
| FMRP targets | 813 | 4.97E-09 | 7.19E-06 | 4.76E-07 | - |
| Synaptic genes | 1,046 | 4.36E-07 | 5.55E-05 | - | 4.43E-05 |
| Brain-expressed genes | 9,695 | 6.13E-19 | - | 6.68E-17 | 7.55E-16 |

Gene-level analysis performed via MAGMA of relevant gene-sets: i) categories under this study: genes containing canonical CPE, genes bound by CPEB1, CPEB3, and CPEB4; ii) genes belonging to categories known to play a substantial role in psychiatry: FMRP interactor targets, specific genes at synapses, and brain-expressed genes. The latter psychiatric-related gene sets were used also to perform a conditional analysis. For each gene-set is shown the number of genes (N genes) and *P*-value of enrichment with SCZ-associated genes after gene-based analysis.

### Decreased inclusion of the neuronal specific microexon of CPEB4 in SCZ

To explore if the splicing alteration of CPEB4 seen in brains of ASD cases also happens in SCZ, we performed vast-tools (Tapial et al., 2017) analysis on the (BA46) cortex RNA-seq data from people with SCZ (n=95) and matched controls (n=75) of the PsychENCODE Consortium BrainGVEX RNA-seq study (PsychEncode-Consortium et al., 2015). After quality control (QC) procedures (see methods), 54 control and 66 SCZ samples met the thresholds for subsequent splicing analyses. Regarding the four CPEB genes, the only skipped exon (SE) event that differed significantly between controls and SCZ was exon 4 of CPEB4 (i.e. the 24 nucleotide microexon) (Supplementary Table 2). Akin to ASD, inclusion of this microexon was significantly reduced in those with SCZ compared with controls. More precisely, in average, 64.1% of the CPEB4 transcripts contained the microexon in the control samples versus 60.4% in the SCZ samples, resulting in a significant percent spliced in index difference (ΔPSI) = -3.63 (*P*=0.023) (Fig. 1A).

**Figure 1:**
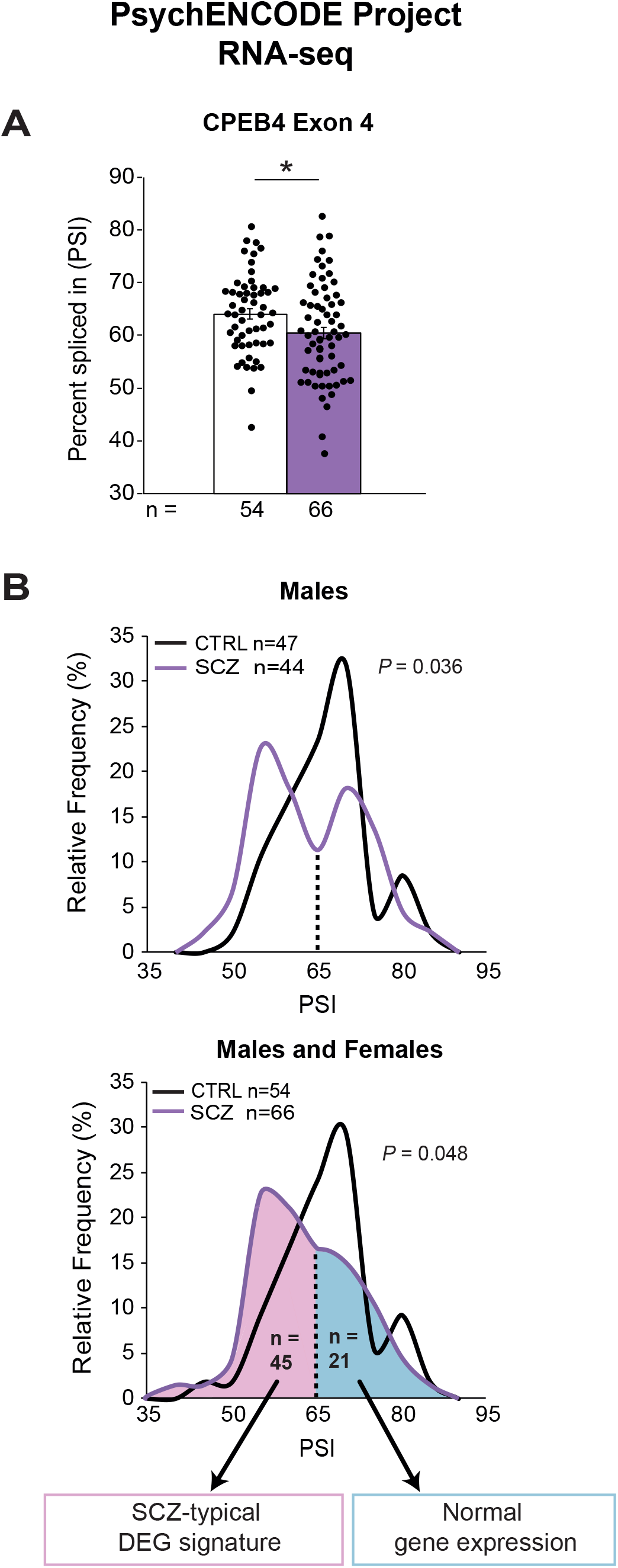
Decreased inclusion of CPEB4 exon 4 in SCZ. Vast-tools splicing analysis of PsychENCODE BrainGVEX data. A) Representation of CPEB4 exon 4 PSI and B) its relative frequency (%) distributions in controls and SCZ patients. Males (upper panel), males and females (lower panel). In the lower panel, SCZ individuals with CPEB4 exon 4 PSI<65% are highlighted in pink and those with CPEB4 exon 4 PSI>65% in light blue. Analysis of differentially expressed genes (DEGs) was performed comparing each group with controls. A) Student’s T-test. B) Two-sample Kolmogorov-Smirnov test (upper panel) and Student’s t-test followed by Benjamini-Hochberg correction for multiple comparison (bottom panel). \**P*<0.05. Data are mean with ±SEM.

### Marked exon 4 skipping correlates with typical SCZ-transcriptomic signature

When we analyzed the relative frequencies of the percentages of inclusion of CPEB4 exon 4, we noticed that the distributions significantly differed between control and SZC samples (*P*=0.048) (Fig. 1B, lower panel). When restricting the analysis to only male samples, two peaks could be clearly observed in the SCZ distribution, the major peak corresponded to a value of percent spliced in index (PSI)≈55, while the minor peak corresponded to PSI≈70, the latter matching with the mode value in the distribution of control samples (Fig. 1B, upper panel). Therefore, in terms of inclusion of exon 4 of CPEB4, the observed distribution in SCZ seem to represent two subpopulations, one that resembles controls, and one whose peak value differs from that in controls with ΔPSI=-15. To further explore whether the two parts of the distribution may represent two different subpopulations, we interrogated global differences in transcript levels with respect to controls. In view of the inflections of SCZ distributions at PSI=65 (Fig. 1B), we stratified the 66 SCZ samples analyzed in Fig. 1a into two pools based on this cut-off. Interestingly, gene expression at the PSI>65-SCZ subpopulation (n=21) was similar that of control samples (Supplementary Table 3). In contrast, the PSI<65-SCZ subpopulation (n=45) showed 771 differentially expressed genes (DEG), 492 upregulated and 279 downregulated (Supplementary Table 3). Remarkably, the DEG signature in the PSI<65-SCZ subpopulation, particularly the downregulated genes, is highly concordant with DEG signatures previously reported for dorsolateral prefrontal cortex (DLPFC) of SCZ subjects^32-33^. More precisely, the representation factor (see methods) was 9.3 (*P*<1.17×10^−7^) regarding RNA-seq based signatures (Yang et al., 2020) and 12.7 (*P*<1.94×10^−6^) regarding gene-chip analysis (Hashimoto et al., 2008), with marked expression deficits in GABA neurotransmission-related transcript such as GAD1 (GAD67) or the neuropeptides somatostatin (SST), neuropeptide Y (NPY) and cholecystokinin (CCK) (Supplementary Table 4). These results indicate that the SCZ individuals seem to segregate into two subpopulations, one that matches controls in terms of both CPEB4 exon 4 inclusion (PSI≈70) and global gene expression, and the other with lower CPEB4 exon 4 inclusion (PSI≈ 55) and a paradigmatic SCZ DEG signature. This suggests interrelated alterations of transcription and of CPEB4-dependent translational regulation in SCZ.

### CPEB4 mis-splicing is not observed in antipsychotic-treated patients

There is evidence of antipsychotic medication correlating with diminished alteration of protein expression in post-mortem brains from individuals with schizophrenia (Chan et al., 2011). This led us to speculate that the two SCZ subpopulations arbitrarily delimited with the PSI=65 cut-off value might correlate with a different degree of exposure to antipsychotic drugs (APDs).

Interestingly, the PsychENCODE database metadata provide “Lifetime Antipsychotics” index value for 45 of the 66 SCZ analyzed samples and, when we analyzed the mean Lifetime APDs index values for each group, we found that it was significantly higher in the PSI>65-SCZ (n=15) subpopulation compared to PSI<65-SCZ (n=30) subpopulation (95,237±20,179 vs. 42,770±7,786; *P*=0.048) (Fig. 2A). This observation suggests that marked alteration in CPEB4 exon 4 inclusion might be specific to SCZ individuals with lower exposure to antipsychotic medication.

**Figure 2:**
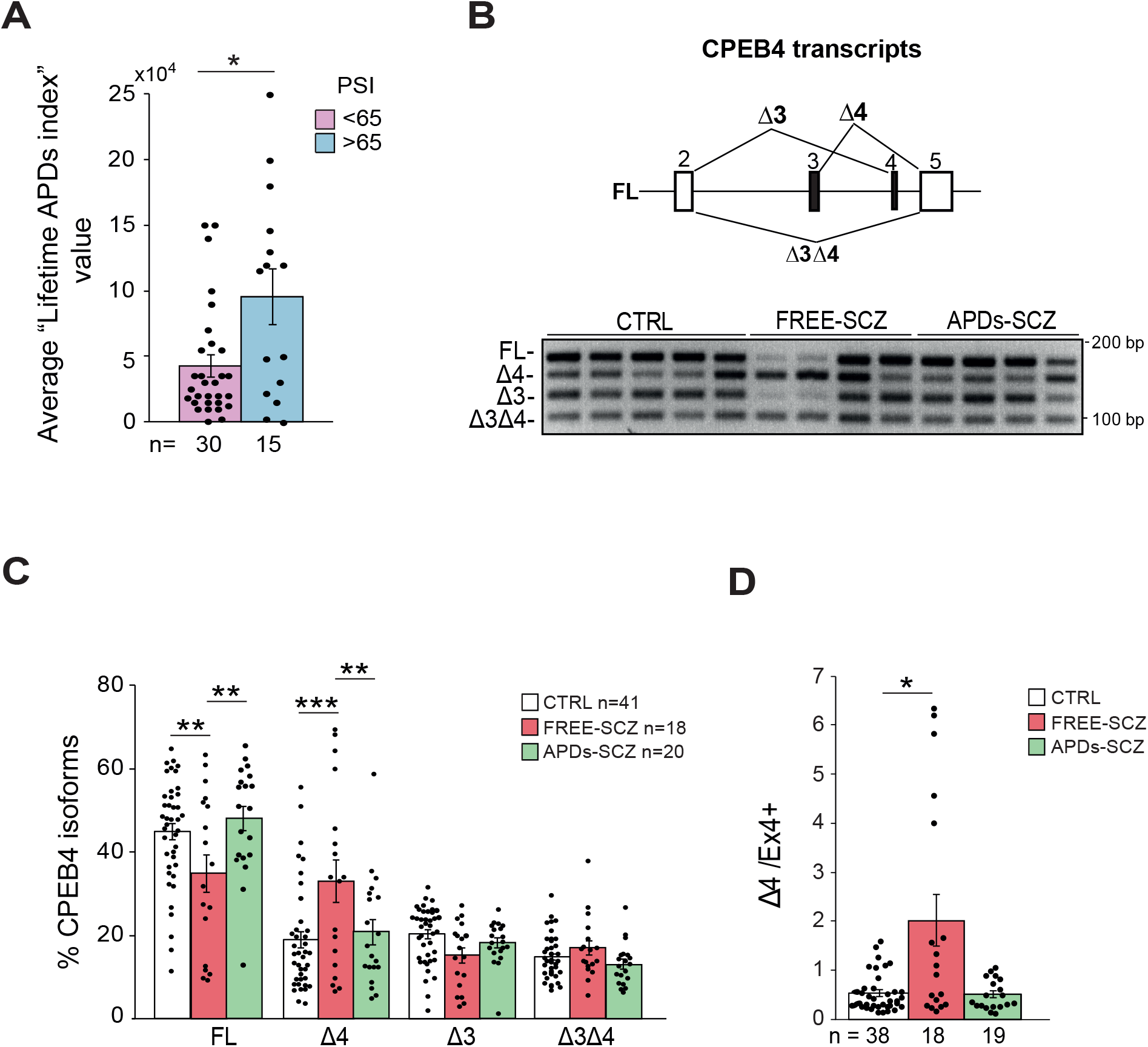
CPEB4 mis-splicing selectively occurs in antipsychotic-free SCZ brains. A) PsychENCODE BrainGVEX project(PsychEncode-Consortium et al., 2015) provides the value of “Lifetime antipsychotic drugs consumption index” for 45 of the SCZ samples used in our RNA-seq analysis. The graph shows the values of this parameter in the SCZ individuals with CPEB4 exon 4 PSI<65 and in those with CPEB4 exon 4 PSI>65. B) At the top, diagram of the four splicing variants of CPEB4. At the bottom, representative RT–PCR analysis of brain tissue from controls and SCZ subjects negative (FREE-SCZ) or positive to antipsychotic drugs (APDs-SCZ) at the moment of death, with C) the corresponding quantification of the percentage of CPEB4 isoforms. D) ΔEx4/Ex4+ ratio. A) Mann-Whitney test C) Two-Way ANOVA test with Bonferroni Correction D) Kruskal-Wallis test with Dunn’s multiple correction. \**P*<0.05, \*\**P*<0.01, \*\*\**P*<0.001. Data are mean with ±SEM.

To confirm in an independent cohort of samples the decreased usage of exon 4 in SCZ observed through RNA-seq analysis and to settle or discard the correlation with antipsychotic medication, we decided to perform RT-PCR analysis in SCZ postmortem DLPFC samples from the Basque Institute of Legal Medicine and the NIH NeuroBioBank (CTRL n=57 and SCZ n=42) in which a complete toxicological examination was performed by mass spectrometry to detect the presence of antipsychotics, as well as mood stabilizers, cotinine, antidepressants and benzodiazepines, at the time of death. Controls positive for any substance, as well as samples with low RNA quality, were excluded from the analysis (see methods). Since exon 3 (57 nucleotides) of CPEB4 is also alternatively spliced (Fig. 2B), there are four CPEB4 isoforms that can be detected with PCR primers hybridizing to exons 2 and 5, two transcripts that include exon 4 (full-length (FL) and CPEB4Δ3) and two lacking it (CPEB4Δ4 and CPEB4Δ3Δ4) (Fig. 2B). In control samples, the FL-CPEB4 transcript predominates (Fig. 2B-C) and, as expected from the RNA-seq data, FL-CPEB4 decreases in samples from SCZ individuals that were free of antipsychotics at the time of death (FREE-SCZ) (Fig. 2B-C). As expected, patients of the FREE-SCZ sample show an increase of CPEB4Δ4 compared to controls (Fig. 2B-C), with some of the FREE-SCZ samples showing a completely opposite pattern of isoforms respect to controls, as CPEB4Δ4 and CPEB4Δ3Δ4 clearly predominate (Fig. 2B). As suggested by both the RNA-seq and Lifetime APDs index analyses, SCZ individuals under APDs medication at the time of death (APDs-SCZ) significantly differed from FREE-SCZ samples (Fig. 2C), showing FL-CPEB4 and CPEB4Δ4 isoform levels to be indistinguishable from those in controls (Fig. 2C). Consequently, the ΔEx4/Ex4+ isoform ratio is markedly increased in FREE-SCZ samples, while unaltered in APDs-SCZ samples (Fig. 2D). Together, these results demonstrate that individuals with SCZ show a shift in the ratio of exon 4-dependent CPEB4 transcripts, selectively in the absence of antipsychotic medication.

### Decreased protein levels of CPEB4 target SCZ genes in antipsychotic-free SCZ brains

As mentioned, the increase of CPEB4Δ4 transcript in brains of idiopathic ASD patients correlates with concerted decreased protein expression of multiple ASD risk genes that are targets of CPEB4 (Parras et al., 2018). Since we have found an enrichment of CPE-containing and CPEB4-binding transcripts among SCZ susceptibility genes (Table 1), we hypothesized that the observed increase of CPEB4Δ4 in DLPFC of antipsychotic-free SCZ cases might result in decreased protein levels of multiple CPEB4-target SCZ risk genes. To test this by Western blot, we identified, among the top 5% SCZ risk genes with the most significant *P*-values in the MAGMA analysis, those i) that are brain expressed, ii) that are binders of CPEB4 (but not of CPEB1, to maximize chances of detecting CPEB4Δ4-associated changes), iii) that show evolutionary conserved presence in the 3’UTR of canonical CPE sequences in human and mice, and iv) whose transcript levels do not vary between control and SCZ samples in our analysis of PsychEncode RNA-seq data (in order to be able to show effects on translation independent of mRNA). This resulted in a short list of 43 top candidate genes (Supplementary Table 5) and we assayed antibodies for 14 of them in Western blot analyses. Seven of these antibodies yielded protein signal at the predicted molecular weight: *BCL11A, CACNB2, CNTN4, CTNND1, OSBPL3, STAG1* and *TCF4* (Supplementary Table 5). Interestingly, the protein levels of *BCL11A, CTNND1, OSBPL3, STAG1* and *TCF4* were reduced in the FREE-SCZ samples, but not in the APDs-SCZ samples (Fig. 3). Furthermore, through a manual curation we identified, among the 5% SCZ risk genes with the most significant *P*-values in the MAGMA analysis, four genes (*GABBR2, HCN1, NEK1* and *SOX5*) that could not have been detected as CPEB4 binders in the CPEB4-RIP experiment performed on striatal tissue (Parras et al., 2018) because they show minimal expression in striatum (according to our published RNA-seq datasets (Elorza et al., 2021)) but that are potentially interesting to this study because they are expressed in prefrontal cortex (according to our analysis of PsychEncode RNA-seq data) and they contain numerous CPEs (at least 4 in both human and mouse). Interestingly, among them, NEK1 has been reported to be translationally regulated by CPEBs in a CPEB4-related paper (Pascual et al., 2020) and we found that NEK1 protein levels are also decreased in the FREE-SCZ samples, but not in the APDs-SCZ samples (Fig. 3). Together, these results indicate that the increase of transcripts excluding exon 4 observed in postmortem DLPFC samples of SCZ cases free of antipsychotic medication correlates with decreased protein levels -despite unaltered transcript levels-, of multiple SCZ-associated genes that are targets of CPEB4.

**Figure 3:**
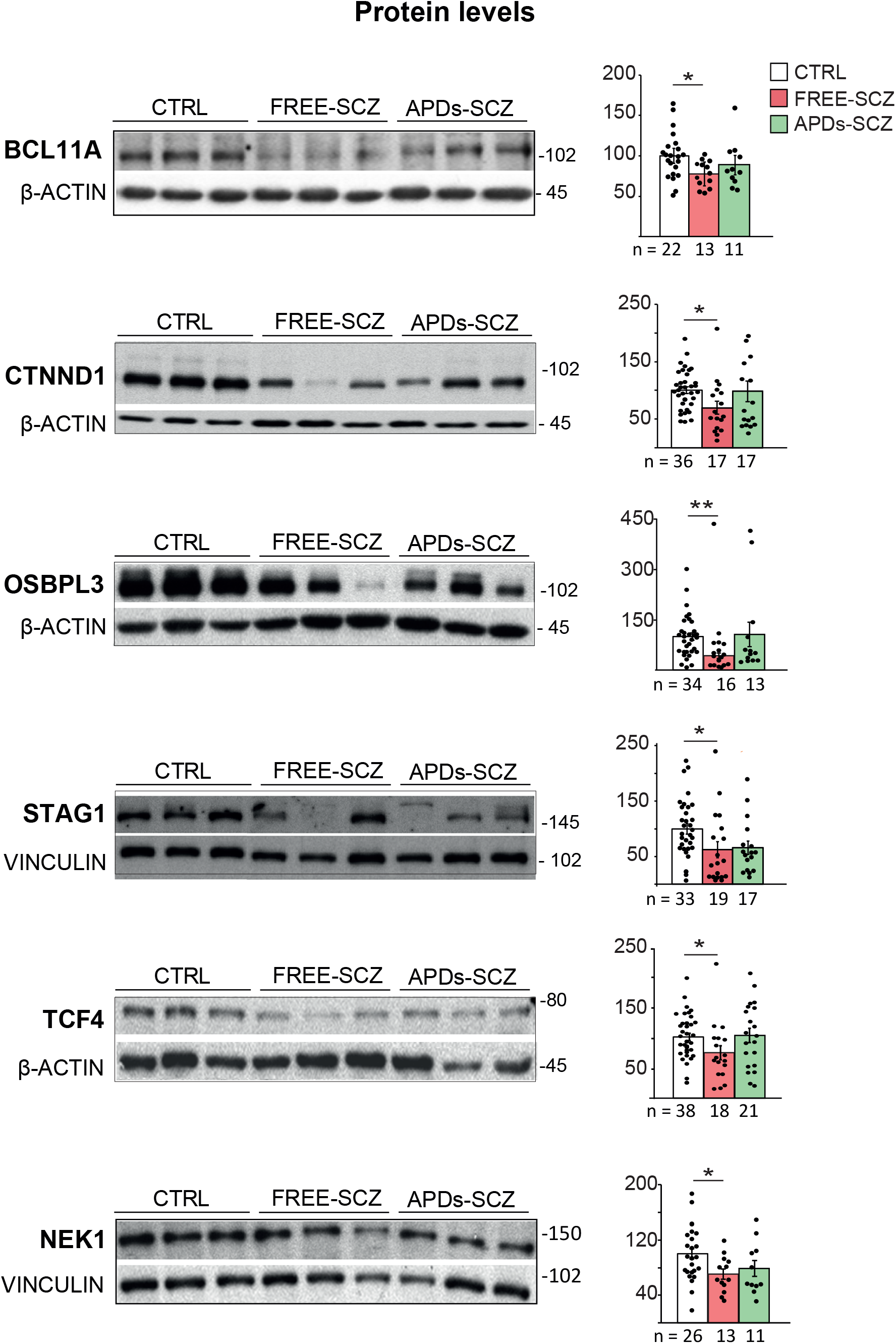
Decreased protein levels of the SCZ risk genes in FREE-SCZ brain samples. TCF4, STAG1, CTNND1, OSBPL3 and NEK1 protein levels in control, FREE-SCZ and APDs-SCZ samples. Kruskal-Wallis test with Dunn’s multiple comparison test or One-Way ANOVA test. \**P*<0.05, \*\**P*<0.01. Data are mean with ±SEM.

### Decreased protein levels of CPE-containing and CPEB4-target SCZ genes in CPEB4Δ4 overexpressing mice

Thus far, we have provided correlative evidence of the role of CPEB4 mis-splicing on expression of SCZ risk genes. To test *in vivo* whether an increase of CPEB4Δ4 transcript suffices to induce concerted decreased protein levels of multiple SCZ risk genes, we leveraged of a previously generated transgenic mouse line that allows conditional overexpression of CPEB4Δ4 in forebrain neurons at different time points and levels (through a tetracycline-system controlled transactivator) (Parras et al., 2018). Since strong CPEB4Δ4 overexpression starting at embryonic stages leads to consistent ASD like phenotypes already evident in pups (Parras et al., 2018), in the context of this SCZ related study, we decided to use a transactivator mouse line with milder expression starting postnatally (PN), to generate Tg-PN-CPEB4Δ4 mice. Young (6 weeks) Tg-PN-CPEB4Δ4 mice show detectable overexpression of CPEB4Δ4 transcript (Fig. 4A) and when we performed Western blot analysis of the CPE-containing SCZ genes that we found decreased in the human FREE-SCZ samples, we also observed decreased protein levels of BCL11A, OSBPL3, TCF4 and NEK1 in Tg-PN-CPEB4Δ4 mice (Fig. 4B). This therefore demonstrates that modest overexpression of CPEB4Δ4 suffices to cause concerted under-expression of multiple proteins encoded by genes associated to SCZ.

**Figure 4:**
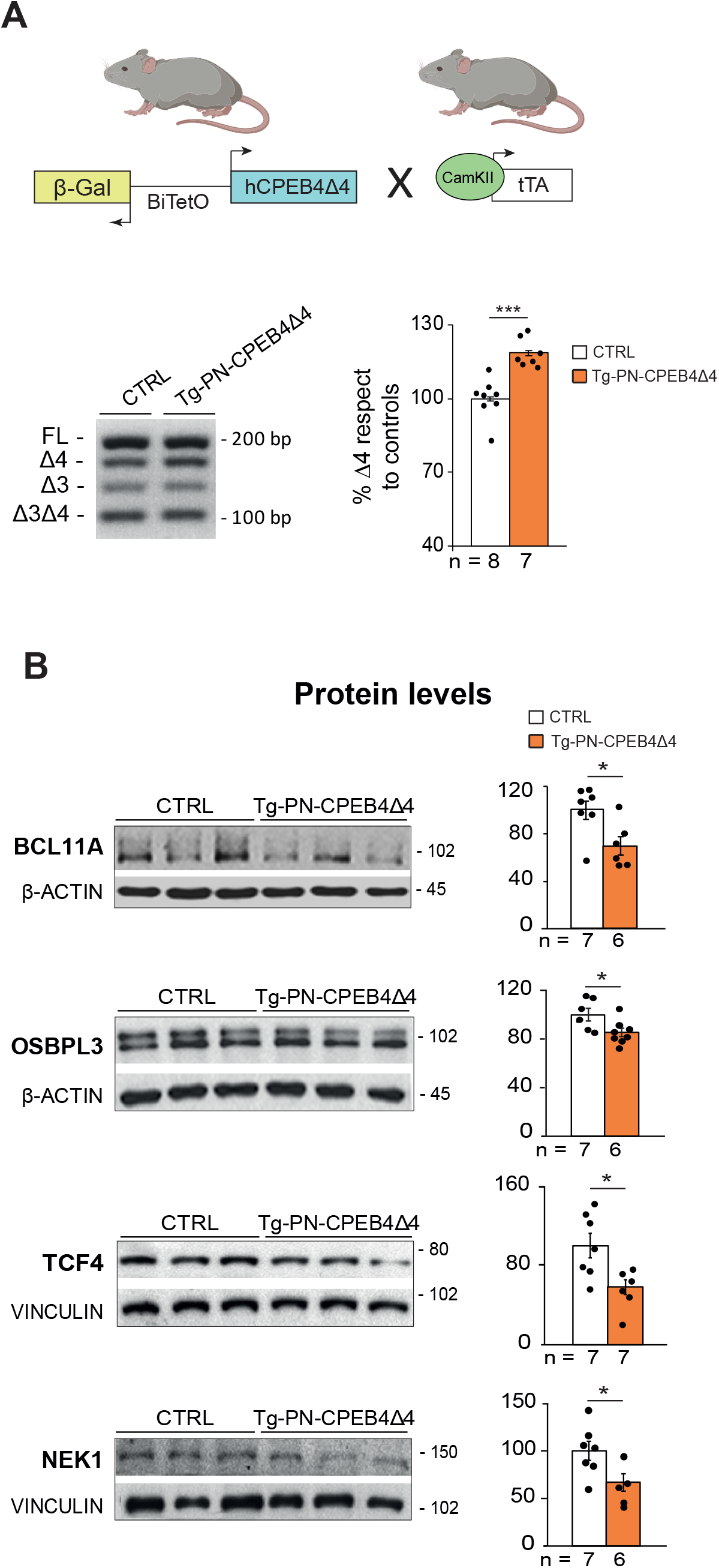
Transgenic overexpression of CPEB4Δ4 isoform in mouse brains suffices to induce decreased TCF4, OSBPL3 and NEK1 protein levels. A) Scheme of the transgenes in the Tg-PN-CPEB4Δ4 mouse model (top). Representative RT-PCR of CPEB4Δ4 isoform in control and Tg-PN-CPEB4Δ4 mouse brain tissue with corresponding quantification (bottom). B) TCF4, OSBPL3 and NEK1 protein levels in control and Tg-PN-CPEB4Δ4 mouse brains. A) and B) Student’s t-test; \**P*<0.05, \*\**P*<0.01, \*\*\**P*<0.001. Data are mean ±SEM.

### CPEB4Δ4 overexpressing mice show SCZ-linked behaviors

To gain insight into whether the overexpression of CPEB4Δ4 with subsequent mis-expression of SCZ susceptibility genes lead to SCZ-like behaviors in mice, we analyzed the pre-pulse inhibition (PPI) of the startle response (SR) in Tg-PN-CPEB4Δ4 mice. As shown in Fig. 5A, SR response is normal in Tg-PN-CPEB4Δ4 mice at all frequencies, thus ruling out any hearing impairment and, importantly, Tg-PN-CPEB4Δ4 mice showed impaired PPI of the SR, therefore confirming a SCZ-like rodent phenotype. Tg-PN-CPEB4Δ4 mice also showed other altered behaviors frequent in animal models of SCZ, such as increased grooming and decreased social interaction (Fig. 5B-C). These data therefore demonstrate that *in vivo* overexpression of CPEB4Δ4 suffices to induce SCZ-related behavioral alterations of mouse models.

**Figure 5:**
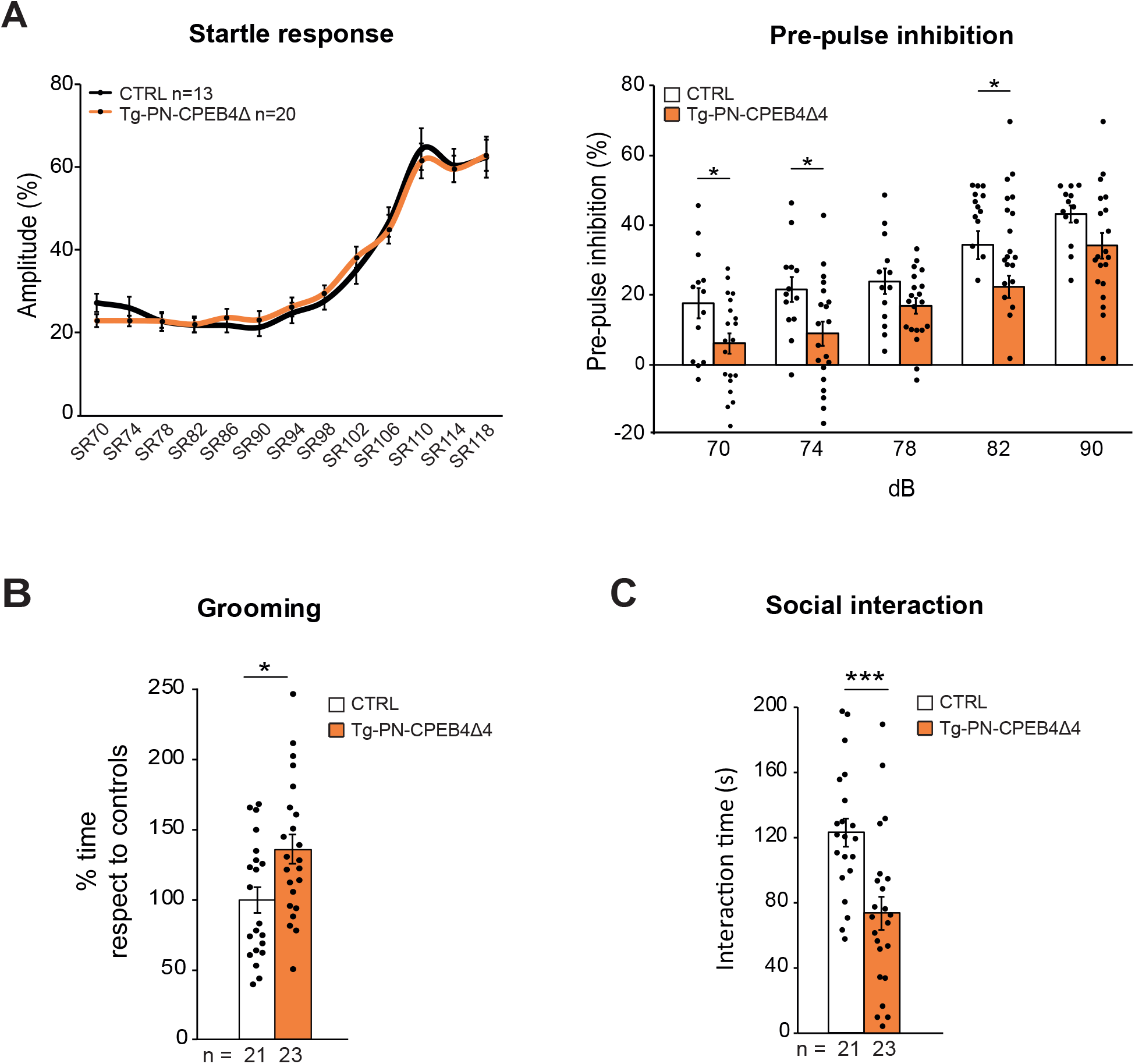
Tg-PN-CPEB4Δ4 mice show schizophrenia-linked rodent behaviors. A) Representation of the amplitude of acoustic startle response corresponding to acoustic stimuli of increasing decibels (from 70 to 118 dB) in control and Tg-CPEB4Δ4 mice (left); quantification of the pre-pulse inhibition (PPI) of the acoustic startle response at 70, 74, 78, 82 and 90 Db (right). B) Grooming time during a 5 min trial. C) Interaction time with an unfamiliar mouse in social interaction test. A)-B)-C) Student’s t-test; \**P*<0.05, \*\**P*<0.01, \*\*\**P*<0.001. Data are mean with ±SEM.

## Discussion

In this study we show altered splicing of *CPEB4* in DLPFC of SCZ patients. CPEB4 belongs to a family of RNA-binding proteins that regulate the translation of specific mRNAs containing CPE sequences in their 3’ untranslated regions (UTRs), targeting about 40% of transcripts. CPEBs play important roles in development and neuronal plasticity (Ivshina et al., 2014) and we have previously demonstrated CPEB4 mis-splicing in ASD brains leading to concerted mis-expression of a plethora of high confidence ASD risk genes (Parras et al., 2018). Using data from the largest GWAS meta-analysis in SCZ (Trubetskoy et al., 2022), we find that both CPE-harboring and CPEB4-binder gene subsets are significantly enriched in SCZ associated genes. Through RT-PCR and Western blot analyses on postmortem DLPFC tissue, we further characterize the CPEB4 transcript isoform switch in SCZ, which occurs specifically in antipsychotic medication-free individuals. Furthermore, the imbalance in CPEB4 isoforms correlates with diminished protein levels of top SCZ associated genes being either CPE-harboring and/or CPEB4-binder. Finally, we demonstrate that mild overexpression of CPEB4Δ4 in transgenic mice (to mimic the CPEB4 transcript isoform imbalance observed in SCZ brains) suffices to cause decreased protein levels of target SCZ-risk genes and to induce SCZ-associated behaviors found in mouse models of the disease.

Since our mouse genetics data demonstrate a causal role of CPEB4 mis-splicing on SCZ-associated gene expression and behaviors, our study pinpoints CPEB4 mis-splicing as a potential novel target for therapeutic intervention in SCZ. Interestingly, the splicing alteration is not observed in patients that were on antipsychotics medication at the time of death, suggesting that the current medication that improves symptoms in SCZ patients, also results in normalization of CPEB4 splicing. Recently, splicing modifying therapies, particularly antisense oligonucleotides (ASOs) have reached the clinic for chronic neural and neuromuscular conditions such as spinal muscular atrophy and Duchenne muscular dystrophy (Rinaldi and Wood, 2018). It is therefore conceivable that CPEB4 splicing-modifying ASOs, if administered in combination with antipsychotics, might improve the efficacy of the treatment.

Broad splicing alteration has been suggested to play a role in SCZ (Zhang et al., 2022) and it would be interesting to disentangle the molecular mechanism underneath that global aberrant RNA splicing in SCZ, and behind CPEB4 mis-splicing in particular. This might be useful to design interventions with small molecule drugs, as has been proposed for cancer (Dong and Chen, 2020). Since the CPEB4 splicing alteration is not seen in patients taking antipsychotics and the SCZ associated transcriptomic signature is observed only in the RNAseq-datasets of patients that show aberrant inclusion of CPEB4 exon 4 (PSI<65) -which in turn correlate with lower Lifetime antipsychotics index-, screening the RNA-seq data for splicing factors mis-expressed in SCZ PSI<65 but not in SCZ PSI>65 might be informative to identify potential splicing factors responsible for both broad and CPEB4-specific mis-splicing in SCZ.

Regarding the mechanism leading to altered splicing machinery selectively in individuals not receiving antipsychotic therapy, markers of GABAergic neurons are abundant among top downregulated genes in SCZ PSI<65 samples (while SCZ PSI>65 samples are indistinguishable from controls in terms of gene expression). The latter fits the excitation/inhibition (E/I) imbalance model of SCZ and ASD etiology (Gao and Penzes, 2015). It is therefore conceivable that E/I imbalance in SCZ results in altered gene expression -including splicing factors governing the inclusion of CPEB4 microexon-, with antipsychotics attenuating the E/I imbalance (Llado-Pelfort et al., 2016; Smucny et al., 2022), thus counteracting abnormal expression of the relevant splicing factors and, hence, CPEB4 mis-splicing. In good agreement, it has been reported that inclusion of the program of ASD-associated microexons is neuronal activity-dependent (Quesnel-Vallieres et al., 2016).

This study further supports that, within the ‘neurodevelopmental model’ of psychiatric diseases (Birnbaum and Weinberger, 2017), ASD and SCZ share common molecular mechanisms as defective inclusion of CPEB4 microexon has previously been reported in ASD, with subsequent diminished protein levels of ASD-risk genes (Parras et al., 2018). A common assumption in the neurodevelopmental model is that the etiological process is relatively subtle in SCZ, such that it could be compensated for early in life, while it is stronger in ASD, leading to greater developmental pathology or disruption of neural functions that is not compensable early in life(Birnbaum and Weinberger, 2017; Owen and O’Donovan, 2017). In line with this model, moderate overexpression of the CPEB4Δ4 transcript isoform starting postnatally in Tg-PN-CPEB4Δ4 mice results in SCZ-relevant phenotypes, like decreased PPI and social interaction, in the absence of overt visible phenotypes. On the contrary, robust overexpression of CPEB4Δ4 transcript stating at embryonic stages results in evident phenotypes such as stereotypic running in the periphery of the mouse home cage and hydrocephalous, apart from more subtle ASD-associated behaviors (Parras et al., 2018). Interestingly, although decreased social interaction is common in mice with moderate and strong expression of CPEB4Δ4, diminished PPI is only found in Tg-PN-CPEB4Δ4 mice, as mice with prenatally-starting strong CPEB4Δ4 overexpression show a different behavior in the PPI test consisting on no effect of high intensity prepulses and even a tendency to increased PPI at low intensity prepulses (data not shown).

In summary, our study unveils CPEB4 mis-splicing -consisting in reduced inclusion of a neuronal microexon- and concomitant decreased protein levels of CPEB4-target SCZ genes as a new molecular mechanism in SCZ and further support the etiological parallelism between SCZ and ASD.

## Materials / Subjects and Methods

### CPEBs-related genes enrichment analysis among SCZ risk genes

To establish a relationship between SCZ genes and CPEBs, we carried out gene-set enrichment analyses through MAGMA, a bioinformatics tool for gene-level analyses of GWAS data sets (de Leeuw et al., 2015). European and East Asian SCZ GWAS summary statistics were obtained from the latest public multi-stage study of the Psychiatric Genomics Consortium (Trubetskoy et al., 2022), and processed through a gene-level analysis in MAGMA as described therein. Briefly, this involved computing gene-wide mean *P*-values with each ancestry-specific subsample, which were then meta-analysed leading to combined gene-level data from 67,390 SCZ cases and 94,015 controls. For these analyses, gene boundaries were retrieved from Refseq (GRCh38.p13; last updated on 23-11-2020), lifted over to GRCh37 coordinates and expanded by 35kb/10kb upstream/downstream flanking regions to encompass potential regulatory elements (Maston et al., 2006). Regarding CPE-containing genes in humans, we considered a previously published database (Pique et al., 2008). CPEB1 and CPEB4 gene lists were derived from an RNA immunoprecipitation (RIP) study performed in mouse striatum(Parras et al., 2018). A list of CPEB3 binders has been recently obtained through a RIP experiment performed in mouse cortical tissue (Lu et al., 2021). Covariate gene sets used to carry out conditional analyses included (i) targets of the FMRP regulon (Darnell et al., 2011), (ii) genes with significant expression in the brain (Fagerberg et al., 2014); as processed by (Genovese et al., 2016) and (iii) genes specifically involved in synaptic development and function (Koopmans et al., 2019).

### RNA-seq data analysis

To explore the alternatively spliced events of CPEBs in SCZ patients, we examined the data in the BrainGVEX RNA-seq study^31^. Such study comprises well-characterized human post-mortem prefrontal (BA46) cortex samples from SCZ subjects (n=95) and matched controls (n=75), originating from the Stanley Medical Research Institute (SMRI). Fastq files were downloaded from the PsychENCODE Consortium (PEC) Knowledge Portal repository (https://psychencode.synapse.org/). For quality control analysis, we run FastQC (v0.11.9). Then, the four parameters i) *per base GC content*, ii) *per sequence GC content*, iii) *sequence duplication level* and iv) *overrepresented sequences* were monitored for each strand (forward and reverse), thus resulting in a total of 8 tests. Finally, only samples that overall pass at least four of the latter were considered. Following the quality filter, a total of n=54 controls and n=66 SCZ samples got access to the final step of the analysis pipeline. Reads were aligned to human hg19 reference genome and counts and per cent spliced in (PSI) values were computed through complete Vertebrate Alternative Splicing and Transcription Tools (vast-tools) v.2.5.1 program (Irimia et al., 2014; Tapial et al., 2017). Regarding CPEB4 Ex4 PSI values relative frequency distribution, a bin width of 10 has been considered. Differentially expressed genes between CTRL and SCZ were obtained calculating the mean of the cRPKM values of each gene and computing the fold change (FC) value. *P*-values obtained with a T-test were corrected by false discovery rare (FDR) method for multiple comparison. Downregulated genes in SCZ with a CPEB4 exon 4 PSI<65 compared to control were obtained with a FDR<1%. In the case of the 100 top genes, a FDR<5% has been chosen. Enrichment analysis of already published SCZ-related DEG signature among our DEGs were performed using http://nemates.org/MA/progs/overlap_stats.html. The representation factor (RF) is the number of overlapping genes divided by the expected number of overlapping genes drawn from two independent groups. A RF>1 indicates more overlap than expected of two independent groups, while a RF<1 indicates less overlap than expected.

### Human brain tissue samples

Brain specimens from the frontal cortex (BA9/BA46) of individuals with SCZ and matched controls used in qRT–PCR and immunoblotting were obtained at autopsies performed in the Basque Institute of Legal Medicine, Bilbao, Spain and by the NIH NeuroBioBank (NBB), Maryland, USA (CTRL n=57 and SCZ n=42). Demographic and clinical feature of individuals are reported in Supplementary Table 6. A toxicological screening for a panel of drugs, including antipsychotics, antidepressant, cotinine and ethanol, was performed on both cohorts by the Central Analysis Service at the University of the Basque Country, Spain. A variety of standard procedures including radioimmunoassay, enzymatic immunoassay, high-performance liquid chromatography and gas chromatography–mass spectrometry have been performed. Following toxicological screening, only controls who resulted negative to any substance were included in the study (n=45), while schizophrenic individuals were divided in a treatment-free group (FREE-SCZ n=21) and a group of antipsychotic treated patients (APDs-SCZ n=21).

### Mice

Conditional transgenic mice carrying a cDNA of human CPEB4 lacking exon 4 (CPEB4Δ4) under control of the inducible TetO promoter were previously generated(Parras et al., 2018) and maintained in a C57BL/6J background. CPEB4Δ4 mice were crossed with a driver mouse line with low expression of the transactivator tTA (Tet-Off) under the CamkII promoter (Low-CamkII-tTA mice) to obtain conditional double transgenic mice with low forebrain neuronal expression of CPEB4Δ4 (Tg-PN-CPEB4Δ4 mice). All mice were housed in the CBMSO animal facility. Mice were housed four per cage with food and water available *ad libitum* and maintained in a temperature-controlled environment on a 12 h–12 h light–dark cycle with light onset at 08:00. Animal housing and maintenance protocols followed local authority guidelines. Animal experiments were performed under protocols approved by the Centro de Biología Molecular Severo Ochoa Institutional Animal Care and Utilization Committee (Comité de Ética de Experimentación Animal del CBM, CEEA-CBM), and Comunidad de Madrid PROEX 247.1/20.

### RNA isolation and cDNA synthesis

Total tissue RNA was extracted from prefrontal cortex (BA9/BA46) of control individuals and matched SCZ using the Maxwell 16 LEV simplyRNA Tissue Kit (Promega, AS1280). Quantification and quality determination of RNA was done on a Nanodrop ND-1000 spectrophotometer and Nanodrop 1000 v.3.7.1 (Thermo Scientific). Only samples with an RNA integrity number (RIN) ≥6 and generating an amplification product were included in our experiments (CTRL n=41; FREE-SCZ n=18 and APDs-SCZ n=20) (Supplementary Table 6). The same conditions were employed in the case of control and Tg-PN-CPEB4Δ4 mice brains. Retrotranscription (RT) reactions were performed using the iScript cDNA Synthesis kit (Bio-Rad, PN170-8891) following the manufacturer’s instructions. In brief, 1,000 ng of total RNA from each sample was combined with 10⍰μl of master mix (includes all necessary reagents along with a mixture of random primers and oligo-dT for priming). The reaction volume was completed up to 40⍰μl with DNase/RNase-free distilled water (Gibco, PN 10977). Thermal conditions consisted of the following steps: 5 min at 25⍰°C; 20 min at 46⍰°C and 1 min at 95⍰°C.

### Quantification of CPEB4 transcript splicing and differential splicing analysis

Specific primers designed in CPEB4 exon 2 (forward, 5⍰-GGACGTTTGACATGCACTCAC-3⍰) and exon 5 (reverse, 5⍰-GAGGTTGATCCCCACGGC-3⍰) able to amplify the four CPEB4 splicing isoforms (full-length, Δ4, Δ3 and Δ3Δ4) in human and mouse brain cDNA were used. PCR amplification protocol was the following: 10 min at 94⍰°C + 33 cycles (30 s at 94⍰°C + 30 s at 58⍰°C + 2 min at 72⍰°C) + 10 min at 72⍰°C. PCR products according with the four CPEB4 isoforms were resolved on 2.2% agarose/GelGreen (Biotium, 41004) gels run at 125 V for 1.5 h. Images were scanned with densitometer (Bio-Rad, GS-900) and quantified with Image Laboratory 5.2 (Bio-Rad). Finally, the percentage of each CPEB4 isoform was calculated.

### Western Blot

Samples from human brain were stored at –80⍰°C and were ground with a mortar in a frozen environment with liquid nitrogen to prevent thawing of the samples, resulting in tissue powder. For mouse, brains were quickly dissected on an ice-cold plate and the different structures stored at –80⍰°C. Human and mouse extracts were prepared by homogenizing the brain areas in ice-cold extraction buffer (20 mM HEPES pH 7.4, 100 mM NaCl, 20 mM NaF, 1% Triton X-100, 1 mM sodium orthovanadate, 1⍰μM okadaic acid, 5 mM sodium pyrophosphate, 30 mM β-glycerophosphate, 5 mM EDTA, protease inhibitors (Complete, Roche, Cat. No 11697498001)). Homogenates were centrifuged at 15,000g for 15 min at 4⍰°C. The resulting supernatant was collected, and protein content determined by Quick Start Bradford kit assay (Bio-Rad, 500-0203). Between 10 and 20⍰μg of total protein was electrophoresed on 10% SDS– polyacrylamide gel, transferred to a nitrocellulose blotting membrane (Amersham Protran 0.45⍰μm, GE Healthcare Life Sciences, 10600002) and blocked in TBS-T (150 mM NaCl, 20 mM Tris–HCl, pH 7.5, 0.1% Tween 20) supplemented with 5% non-fat dry milk. Membranes were incubated overnight at 4⍰°C with the primary antibody in TBS-T supplemented with 5% non-fat dry milk. Antibodies used: mouse anti-ATP2A2 (1:1000, Abcam, ab2861), rabbit anti-ATXN7 (1:1000, Invitrogen, PA1-749), rabbit anti-BCL11A (1:250, Sigma-Aldrich, ABE401), rabbit anti-CACNB2 (1:1000, Santa Cruz Biotechnology, sc-81890), rabbit anti-CNTN4 (1:500, Abcam, ab137107), rabbit anti-CTNND1 (1:1000, Sigma-Aldrich, HPA015955), rabbit anti-ELAVL4 (1:500, Abcam, ab96474), rabbit anti-MEF2C (1:1000, Abcam, ab64644), rabbit anti-NEK1 (1:1000, Thermo Scientific, PA5-15336), rabbit anti-OSBPL3 (1:1000, Santacruz Biotechnology, sc-398326), rabbit anti-PDE4B (1:1000, Abcam, ab14611), rabbit anti-STAG1 (1:1000, kindly provided by Spanish National Cancer Research Center-CNIO)(Remeseiro et al., 2012), rabbit anti-TCF4 (1:1000, Proteintech, 22337-1-AP), rabbit anti-ZEB2 (1:1000, Proteintech, 14026-1-AP), rabbit anti-ZSWIN6 (1:1000, Origene, AP54741PU-N), rabbit anti-VINCULIN (1:10000, Abcam, ab129002) and mouse anti-β-ACTIN (1:25000, Sigma-Aldrich, A2228). Membranes were washed with TBS-T and next incubated with secondary HRP-conjugated anti-mouse IgG (1:2000, DAKO, P0447) or anti-rabbit IgG (1:2000, DAKO, P0448), they were developed using the ECL detection kit (Perkin Elmer, NEL105001EA). Images were scanned with densitometer (Bio-Rad, GS-900) and quantified with Image Laboratory 5.2 (Bio-Rad).

### Mouse Behaviour tests

#### Prepulse inhibition (PPI) of the acoustic startle response test

Startle response curve as well as PPI test were conducted in 10-week-old mice using a commercially available StartFear (Panlab-Harvard Apparatus). This system allows recording and analysis of the signal generated by the animal movement through a high sensitivity weight transducer system. A standard protocol was adapted (Stark et al., 2008). Each mouse has been located in the chamber and during a 5-min acclimation period, while background white noise was continually present. Then, a startle response curve session was performed in order to rule out any impairment in hearing. Startle measures included recordings made every 4 dBs above background (66 dB), up to 118 dB. Each mouse received four times each trial type (40 ms-sound pulses from 70 dB to 118 dB) distributed randomly and separated by 10s-intertrial interval. Response amplitude was considered as the maximum response level recorded during 1 s after the sound pulse. Regarding PPI, trial types, trial type presentation, and background noise levels were performed according to the protocols described previously (Mukai et al., 2004; Stark et al., 2008) with some modifications the following day. In brief, after a 5 min habituation period (66 dB white noise background), eight sets of four different trials distributed randomly with a variable intertrial time (10, 15 or 20 s) were presented to each mouse: trial 1, 40-ms, 120-dB noise burst alone; trials 2 and 3, 120-dB startle stimulus preceded 100 ms earlier by a 20-ms, 70, 74, 78 or 82-dB noise burst (pre-pulse); trial 4, no stimulus, background noise alone (66 dB). As for startle test, response amplitude was considered as the maximum response level recorded during 1 s after the sound pulse. Percent PPI was calculated as 100-[(startle response of acoustic startle from acoustic prepulse and startle stimulus trials/startle response alone trials) × 100].

#### Grooming time and social interaction tests

Grooming and social interaction was examined in 7-week-old mice. The first day (training), mice were allowed to explore for 5 min a three chamber Plexiglas box. This time of habituation was employed to measure the grooming activity. The next day (test), mice were placed in the same box containing two wire cages, one empty and the other with an unknown (gender paired) mouse in it, located in opposite chambers and separated by the empty chamber. Mice were recorded for 10⍰min and the time spent interacting with the unknown mouse was measured.

## Statistical Analysis

Statistical analysis was performed with SPSS 26.0 (SPSS Statistic IBM), GraphPad software (La Jolla, CA, USA) or RStudio 2022.02.2 (Boston, MA, USA). The normality of the data was analysed by Shapiro–Wilk test (n<50) or Kolmogorov–Smirnov test (n > 50). For comparison of two independent groups two-tail unpaired Student’s t-test (data with normal distribution), Mann–Whitney–Wilcoxon or Kolmogorov–Smirnov tests (with non-normal distribution) were performed. For multiple comparisons, data with a normal distribution were analysed by one way-ANOVA or two-way-ANOVA followed by a Tukey’s post hoc test. Statistical significance of non-parametric data for multiple comparisons was determined by Kruskal–Wallis one-way ANOVA. Benjamini-Hochberg correction was applied for multiple testing in RNA-seq analysis. Data are represented as mean⍰±⍰SEM with 95% confidence intervals. Higher or lower points (outliers) are not plotted. A critical value for significance of *P*<0.05 was used throughout the study.

## ACKNOWLEDGEMENTS

RNA-seq data for this publication were obtained from NIMH Repository & Genomics Resource, a centralized national biorepository for genetic studies of psychiatric disorders, specifically, data were generated as part of the PsychENCODE Consortium. For providing brain samples, the authors thank the Basque Institute of Legal Medicine and the Universidad del Pais Vasco (UPV/EHU) as well as Harvard Brain Tissue Resource Center, funded through NIH-NeuroBiobank HHSN-271-2013-00030C, the National Institute of Mental Health (NIMH), National Institute of Neurological Diseases and Stroke (NINDS), National Institute on Aging (NIA), Eunice Kennedy Shriver National Institute of Child Health and Human Development (NICHD), and brain donors and their families for the tissue samples used in these studies.

The authors would like to thank the staff members of the Basque Institute of Legal Medicine for their cooperation in the study. We also thank excellent technical assistance by Miriam Lucas and by the following core facilities: CBMSO-Genomics & Massive Sequencing, CBMSO-Animal Facility and the CETA-CIEMAT computing center.

We thank Dr. M. Irimia (CRG, Barcelona) for advice on AS analysis tools and for critical reading of the manuscript and Dr. Ana Losada (CNIO, Madrid) for providing the anti-STAG1 antibody. This work was supported by CIBERNED-ISCIII collaborative grant PI2018/06-1 to JJL; grants from Spanish Ministry of Economy and Competitiveness/Ministry of Science, Innovation and Universities to JJL: SAF2015-65371-R (MINECO/AEI/FEDER, UE), RTI2018-096322-B-I00 (MCIU/AEI/FEDER,UE) and PID2021-123141OB-I00 (MCIU/AEI/FEDER,UE); to RM: PID2020-119533GB-I00 (MINECO) and to CT: PID2020-114996RB-I00 (MCIU/AEI/FEDER,UE) and RyC2018-024106-I; and grants from Basque Government (IT-1211-19 and 1512-22) to JJM and LFC and from World Cancer Research Fund International (2020_021) to RM. Cardiff University’s work was supported by Medical Research Council Centre (MR/L010305/1), Program (MR/P005748/1), and Project (MR/L011794/1, MC_PC_17212) grants to JTRW, MCOD and MJO. AFP was supported by an Academy of Medical Sciences “Springboard” award (SBF005\1083). IO was hired through European Union’s Horizon 2020 research and innovation program under the Marie Skłodowska-Curie grant agreement (766124).

## AUTHOR CONTRIBUTIONS

I.O. was involved in all assays and data collection, data interpretation and statistical analysis. A.F.P performed the MAGMA analyses. A.P. contributed to experimental design and data interpretation. I.H.H. performed the bioinformatics analyses and contributed to data interpretation. M.S.G. performed bioinformatics analyses and western blotting. S.P. and A.E. contributed to data interpretation, experimental design and discussion. L.F.C. provided patient samples with toxicological data. G.F.M. and E.B. made intellectual contributions, provided reagents and optimized protocols. J.T.R.W and M.C.O’D. made intellectual contributions to and edited the manuscript. C.T. performed bioinformatics analysis and made contribution to the discussion R.M. revised the manuscript and made contribution to the discussion. J.J.M. provided patient samples with toxicological data and made contribution to the discussion. M.J.O. revised the manuscript and made contributions to the discussion. J.J.L. directed the study and designed experiments, drafted and revised the manuscript with input from all authors.

