## Supplementary Table 4 for "Pathogenic mis-splicing of *CPEB4* in schizophrenia"

| **Gene Name** | **FC** |  | **Gene Name** | **FC** |
| --- | --- | --- | --- | --- |
| TNFSF10 | 0,57 |  | BEX5 | 0,76 |
| RERGL | 0,60 |  | NPY | 0,77 |
| PENK | 0,62 |  | DUSP2 | 0,77 |
| DUSP4 | 0,63 |  | NXPH2 | 0,77 |
| TAC1 | 0,65 |  | GIMAP7 | 0,77 |
| CD52 | 0,65 |  | IL34 | 0,77 |
| RPS4Y1 | 0,65 |  | C19orf26 | 0,77 |
| CRH | 0,65 |  | MMD2 | 0,77 |
| PVALB | 0,67 |  | A2M | 0,77 |
| SEMA3G | 0,68 |  | RASL11A | 0,77 |
| LBH | 0,68 |  | FGL2 | 0,78 |
| SST | 0,68 |  | NNAT | 0,78 |
| CXCR4 | 0,68 |  | CRYM | 0,78 |
| OSTN | 0,68 |  | ARL4D | 0,78 |
| ABCG2 | 0,68 |  | HLA-DPB1 | 0,79 |
| HLA-DMB | 0,69 |  | USMG5 | 0,79 |
| TMSB4Y | 0,69 |  | KLF10 | 0,79 |
| KDM5D | 0,69 |  | NUDT14 | 0,79 |
| CX3CR1 | 0,69 |  | ITGAX | 0,79 |
| P2RY12 | 0,69 |  | NPTX2 | 0,79 |
| NEUROD6 | 0,69 |  | ATP5G1 | 0,79 |
| ZFY | 0,71 |  | RLBP1 | 0,79 |
| OLR1 | 0,71 |  | KCNS3 | 0,79 |
| SLC38A5 | 0,71 |  | FABP3 | 0,79 |
| LYVE1 | 0,72 |  | C17orf96 | 0,79 |
| PNOC | 0,72 |  | LGI2 | 0,79 |
| C1orf133 | 0,73 |  | KCNIP3 | 0,79 |
| COX7A2 | 0,73 |  | COX7A1 | 0,80 |
| DUSP1 | 0,73 |  | PROM1 | 0,80 |
| FRZB | 0,73 |  | BDNF | 0,80 |
| OLFML3 | 0,73 |  | ADAP2 | 0,80 |
| MCHR1 | 0,73 |  | UQCRH | 0,80 |
| SMPX | 0,73 |  | NME5 | 0,80 |
| RGS8 | 0,74 |  | HSPA12B | 0,80 |
| GS1-211B7.1 | 0,74 |  | BOLA3 | 0,80 |
| PLA2G5 | 0,74 |  | VGF | 0,80 |
| SPATA2L | 0,74 |  | RASL10A | 0,80 |
| PLD4 | 0,74 |  | FBLN7 | 0,80 |
| CRHBP | 0,74 |  | MGST3 | 0,80 |
| NDUFA4 | 0,74 |  | GAD1 | 0,80 |
| COL5A3 | 0,74 |  | SLC13A5 | 0,80 |
| HSD11B1 | 0,75 |  | TUBA1B | 0,81 |
| VIP | 0,76 |  | UBA7 | 0,81 |
| ETV5 | 0,76 |  | TRPC3 | 0,81 |
| DUSP6 | 0,76 |  | PPEF1 | 0,81 |
| SPEF1 | 0,76 |  | DLK2 | 0,81 |
| IGFBP6 | 0,76 |  | LPAR6 | 0,81 |
| AGPAT9 | 0,76 |  | ACAT2 | 0,81 |
| GIMAP6 | 0,76 |  | CNTN6 | 0,81 |
| LMO2 | 0,76 |  | SCG2 | 0,81 |

**Supplementary Table 4**: Top 100 significantly downregulated genes comparing SCZ with CPEB4 Ex4 PSI<65 (n=45) and CTRL group (n=54). Minimum counts= 1cRPKM; FDR=5%.
